# Identification of a minimal biomarker profile in head-and-neck squamous cell carcinoma tumors

**DOI:** 10.1101/2021.11.12.468359

**Authors:** Laura Sanchez-Diaz, Lola E. Navas, Elisa Suarez-Martinez, Blanca Felipe-Abrio, Ceres Fernández-Rozadilla, Eva M Verdugo-Sivianes, Manuel A. Celis-Romero, Manuel Chaves-Conde, Maria-Dolores Chiara, Yoelsis Garcia-Mayea, Matilde E. LLeonart, Jose Manuel Garcia-Heredia, Sandra Muñoz-Galvan, Angel Carracedo, Juan P. Rodrigo, Amancio Carnero

## Abstract

Although important advances have been made in the knowledge of the molecular mechanisms leading to the development, of head and neck squamous cell carcinoma (HNSCC), only PDL1 is used for the immunotherapy (pemborlizumab) treatment in the first line of metastatic or recurrent disease. There are no other molecular biomarkers currently used in clinical practice. The objective of the study was to identify transcriptional alterations in patients with oral cavity cancer that identify gene networks responsible for resistance to treatment and prognosis. To identify possible targets for the treatment or prevention of these tumors, we screened for changes in transcription of genes that were recurrently altered in patients and that successfully stratify tumoral and non-tumoral samples, as well as patient survival, based on expression levels. The gene panels are primarily related to the cell cycle, DNA damage response, cytokine signaling and the immune system but also to the embryonic stem cell core. Validation of these panels in an independent cohort led to the identification of three non-interconnected genes, WDR66, SERPINH1 and ZNF622, that can predict patient survival and are differentially expressed in 3D cultures from HNSCC primary cell lines. These genes are related to stemness phenotype are transcriptional targets of the pluripotency transcription factors Sox2 and c-Myc. Our results suggest that WDR66, SERPINH1 and ZNF622 con-stitute a minimal signature of stemness transcriptional targets able to predict the prognosis of HNSCC tumors.

**Simple Summary:** The objective of the study was to identify transcriptional alterations in patients with oral cavity cancer to possibly identify gene networks responsible for resistance to treatment and prognosis. We identify bioinformatically gene panels are primarily related to the cell cycle, DNA damage response, cytokine signaling and the immune system but also to the embryonic stem cell core. Validation of these panels in patients independent cohorts led to the identification of three non-interconnected genes, WDR66, SERPINH1 and ZNF622, that can predict patient survival and are differentially expressed in cancer stem cells cultures from HNSCC. These genes are related to stemness phenotype and epithelial-to-mesenchymal transition and are transcriptional targets of the pluripotency transcription factors Sox2 and c-Myc.

## 1. Introduction

Head and neck squamous cell carcinoma (HNSCC) accounts for 4% of all globally diagnosed cancers with more than 640,000 new cases per year, 355,000 annual deaths and a prevalence in human in the range of 2:1 to 4:1 [1, 2]. Alcohol and smoking are common etiological factors [3–5], but exposure to dust from hard alloys, chlorinated solvents [6] and familial genetic predisposition [7] are also risk factors. Moreover, the role of human papillomavirus (HPV) is well established for squamous cell carcinoma of the oropharynx, but its role in laryngeal cancer is uncertain [5, 8, 9]. In addition, these patients are at risk of developing additional tumors due to chronic exposure of the aerodigestive carcinogenic tract: 14% at 5 years, 26% at 10 years and 37% at 15 years [10, 11]. The major prognostic factor of overall survival (OS) is tumor staging, where nodal invasion is more relevant than tumor extension [12–14]. Other prognostic factors include patient comorbidity, ECOG performance status (PS), persistent habits of toxicant consumption, and the location of the primary tumor [15].

Currently, important advances in understanding the molecular mechanisms that lead to the development of head and neck cancer (HNSCC) have been made, and the same complexity in the carcinogenesis of these tumors has been verified in other solid tumors [16]. Knowing the genetic alterations of HNSCC in early stages may help to understand the malignant potential of premalignant lesions and even healthy mucosa in patients exposed to risk factors. Accordingly, detecting possible targets for effective chemoprevention strategies and possible therapeutic targets for patients developing invasive neoplasia would be helpful. However, there are no clinical or molecular biomarkers currently validated in standard practice. Currently, we use PDL1 for the use of immunotherapy (pemborlizumab) in the first line of metastatic or recurrent disease, as response marker [17].

Chronic exposure of the upper aerodigestive tract to carcinogens, such as tobacco, alcohol, human papillomavirus (HPV) infection (mainly in oropharyngeal tumors), Epstein-Barr virus infection in nasopharyngeal tumors, and other carcinogens, results in the progressive accumulation of changes in genes that regulate cell cycle progression, differentiation and mitogenic signaling patterns, angiogenesis and cell death [3, 5, 18]. The sequence of genetic events specific to this process, known as multipath carcinogenesis, is not well defined, and it seems that the accumulation of events (usually affecting 10 or more genes [19, 20], and not necessarily in any particular order) is what determines the histopathological progression from premalignant lesions such as dysplasia to invasive cancers. In head and neck tumors, numerous recurrent chromosomal abnormalities have been reported [20], such as those at 9p21 (containing the tumor suppressor gene CDKN2A encoding p16INK4a and p14ARF), 9q34 (containing the oncogene NOTCH1)[16], 17p13 (containing the tumor suppressor gene tp53), 11p15 (containing the oncogene HRAS), 3q26 (containing the oncogene PIK3A), 3p (containing the tumor suppressor genes FHIT and RASSF1A), 13q21 (containing the tumor suppressor gene BRACA2) and 18q21 (containing the tumor suppressor gene DCC) [21–24]. With these data, the first model of the genetic progression sequence in the carcinogenesis of HNSCC was developed [25]. Recently, inactivation of suppressive oncogenes has been demonstrated through epigenetic mechanisms, such as the hypermethylation of the gene promoters of DCC, RARB (retinoic acid receptor), p16 or MGMT [24]. Lately, the incidence of oropharyngeal squamous cell carcinoma (OPSCC) has increased due to HPV infection, and it presents a better prognosis than those HPV -negative tumours; therefore, it is likely that there exist differences at the molecular level [22, 24].

In this study, we identified molecular alterations in patients with oral cavity squamous cell carcinomas (OCSCC that are likely responsible for resistance to treatment and prognosis. This allowed us to establish a prognostic gene panel to shed light on the tumorigenesis process of HNSCCs and to assess the possibility of using predictive markers of disease behavior as well as identify possible targets for their treatment or prevention. This gene panel contains 200 genes (or a shorter list of 50 genes) that are transcriptionally altered in tumors and are mainly related to the cell cycle, DNA replication, the immune system and, interestingly, they are also related to the embryonic stem cell core. Specific analyses in an independent test cohort showed that the prognostic signature could be narrowed down to 3 genes: WDR66, ZNF622 and SERPINH1. Although they are not interconnected, they may be transcriptional targets for the transcription factors KLF1, Nanog or Sox2. Furthermore, they are more highly expressed in stem cell-enriched 3D cultures. Together, these data suggest that these three genes may be the minimal signature to predict the prognosis and response of HNSCC tumors.

## 2. RESULTS

### 2.1 Identification of biomarkers of poor prognosis in HNSCC tumors

We performed an analysis of 11 public transcriptional datasets (Supplementary Table 1, discovery datasets) to identify genes whose expression may be highly deregulated in oral cavity tumors. We selected those genes that were differentially expressed in more than 50% of the databases (P<0.01), performed a score normalization of their expression fold-change compared with nontumoral tissue and established a score cutoff of more than 2 or more than 3 (either increase or decrease) to obtain two gene sets that we called the ‘long list’ and ‘short list’, respectively (Figure 1A). Genes in the ‘long list’ represent those in the top 5% most deregulated genes, while genes in the ‘short list’ constitute the top 1% most deregulated genes (Figure 1B).

**Figure 1.**
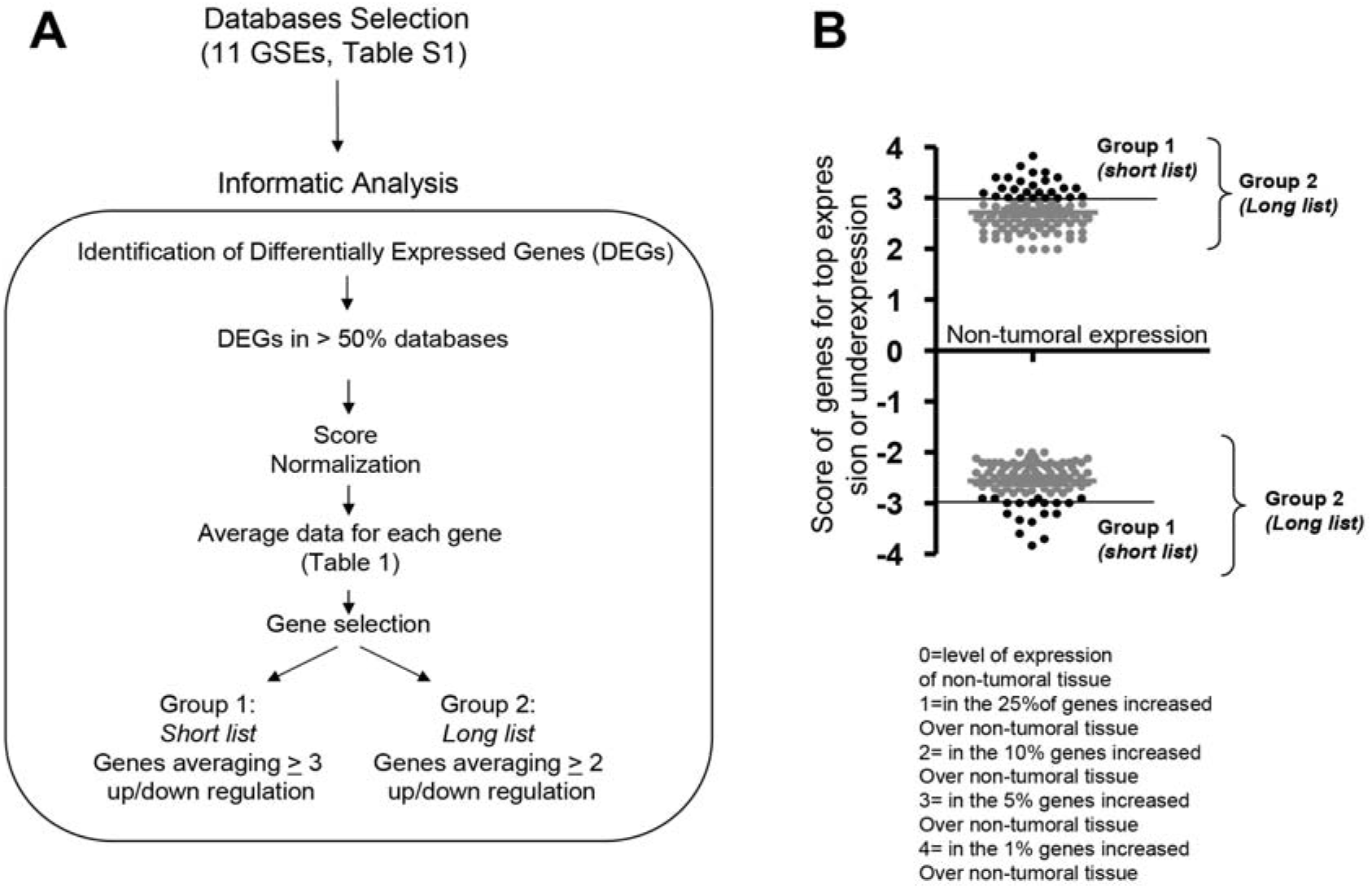
Identification of biomarkers of poor prognosis in oral cavity or HNSCC tumors. (A) Workflow of the analysis performed on 11 public transcriptional datasets to identify genes whose expression may be highly deregulated in oral cavity tumors. (B) Plots of the score of genes for high expression or low expression.

We then analyzed the expression profiles of the two gene sets in patients by hierarchical clustering using the oral cavity, tongue and HNSCC databases. Heatmaps showed that both the ‘long list’ and the ‘short list’ provided a good stratification of patients and control individuals in oral cavity (Figure 2A-B) and tongue squamous cell carcinomas (Supplementary Figure S1) but not in HNSCC tumors (Figure 2C-D and Supplementary Figure S2). These data suggest that the studied genes may be important for OCSCC; therefore, we decided to analyze them fur

**Figure 2.**
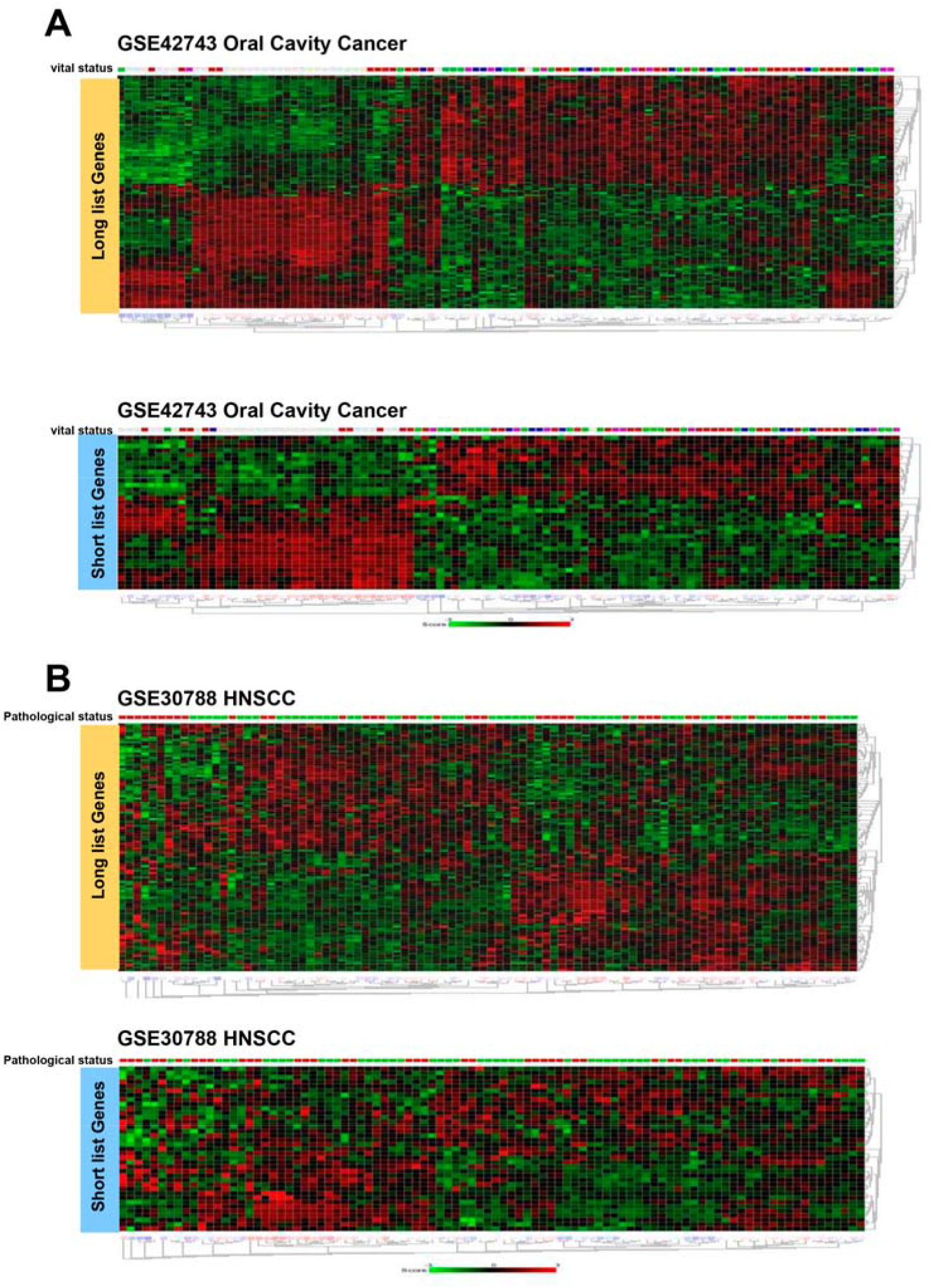
Both long and short gene sets provided a good stratification of patients and control individuals in the oral cavity but were not evident in HNSCC tumors Heatmaps showing the expression z-scores of long list genes (orange) or short list genes (blue) in (A) GSE2743 (oral cavity) and (B) GSE30788 (HNSCC) cancer patient databases.

### 2.2 Network analysis of the gene sets

The gene sets identified above discriminate between tumor and nontumor samples in OCSCC. Therefore, they may be related to one or several specific carcinogenic networks. To perform the network analysis of these gene sets, we first analyzed the physiological or biochemical gene networks to which these genes belonged. We observed that the primary gene sets transcriptionally altered in OCSCC (‘short list’) includes mainly those in the cell cycle, DNA replication and the immune system response Figure 3A). These results were confirmed by Gene Set Enrichment Analysis (GSEA) correlations that showed a clear correlation of the gene sets with Gene Ontology (GO) terms of cell cycle, DNA damage response, cytokine signaling and innate immunity, with a bias towards the upregulated genes (Figure 3B). Notably, we also found GSEA enrichment for the embryonic stem cell core, suggesting a role of this gene set in promoting tumor stemness.

**Figure 3.**
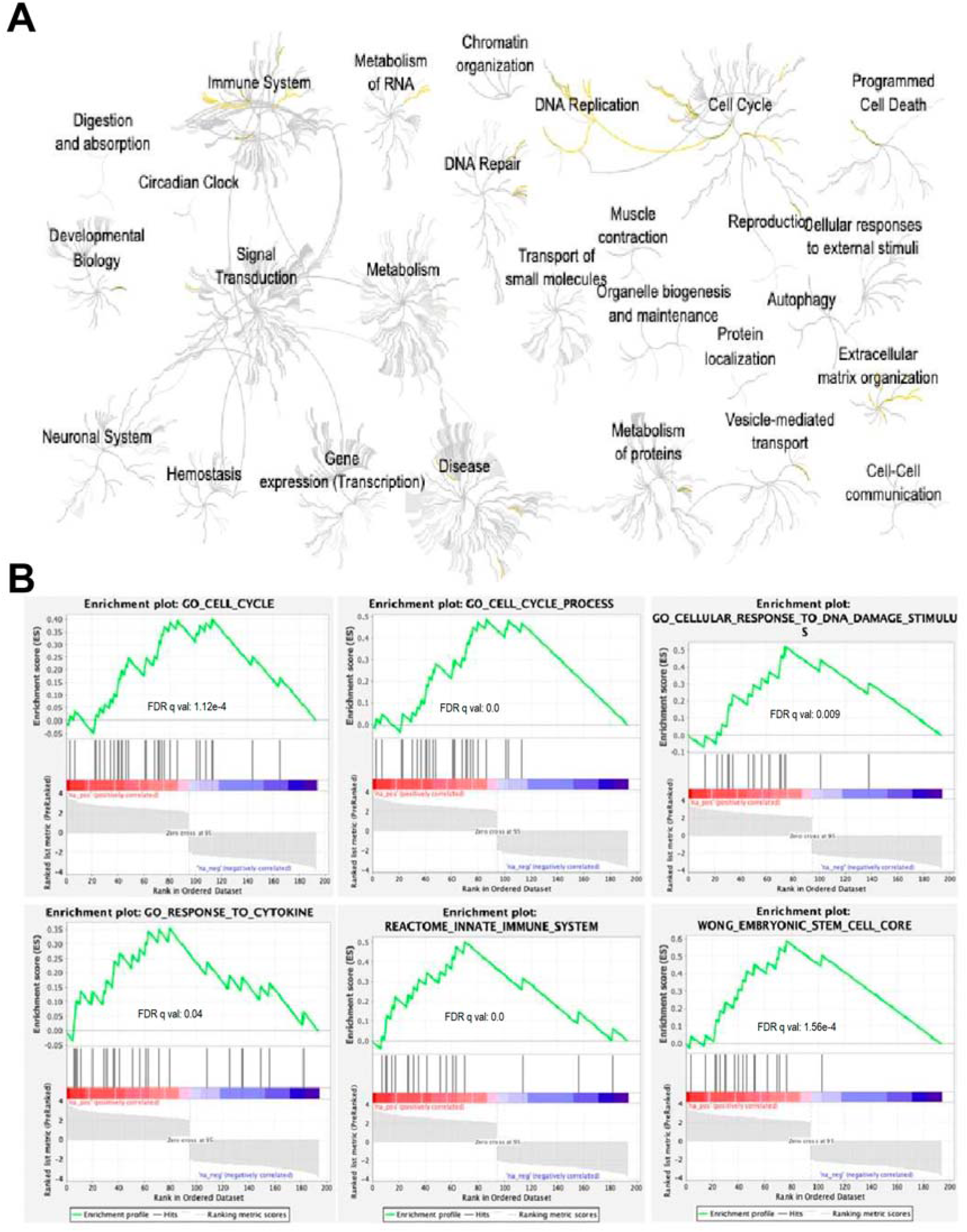
Network analysis of the gene sets. (A) Physiological and biochemical networks related to genes of the “short list” (in yellow). (B) Gene set enrichment analysis (GSEA) correlations of the gene set with Gene Ontology (GO) terms of the cell cycle, DNA damage response, cytokine signaling and innate immunity and cytokine signaling, with a bias towards the upregulated genes.

Further analysis of the enrichment of genes in our gene set discriminating between OCSCC and normal tissue showed clear GSEA enrichment of the transcripts regulated by the factors ASH1L, BARX2, ZSCAN30, SOX6, TCF7 and KLF1 (Figure 4). These genes are related to proliferation regulation and/or gene transcription regulation either by DNA modification, stemness or pluripotency. ASH1l is a histone methyltransferase; BARX2 is a homeobox protein and predicts poor survival in many tumors; ZSCAN30 is a member of the subfamily of zinc finger transcription factors, named zinc finger and SCAN domain containing (ZSCAN) transcription factors, which play several roles in the regulation of cancer progression; and SOX6, TCF7 and KLF1 are related to the pluripotency of stem cells. Finally, to complement our network analysis, we analyzed our gene set for GSEA correlation with pathways in different datasets (Supplementary Figure S3), noting that our gene profile correlated with dividing vs normal quiescent transcription (Graham-CML dataset). We also found a correlation with BRCA1 mutation or Check2 expression (Pujana dataset) and EGFR stimulus (Kobayashy dataset).

**Figure 4.**
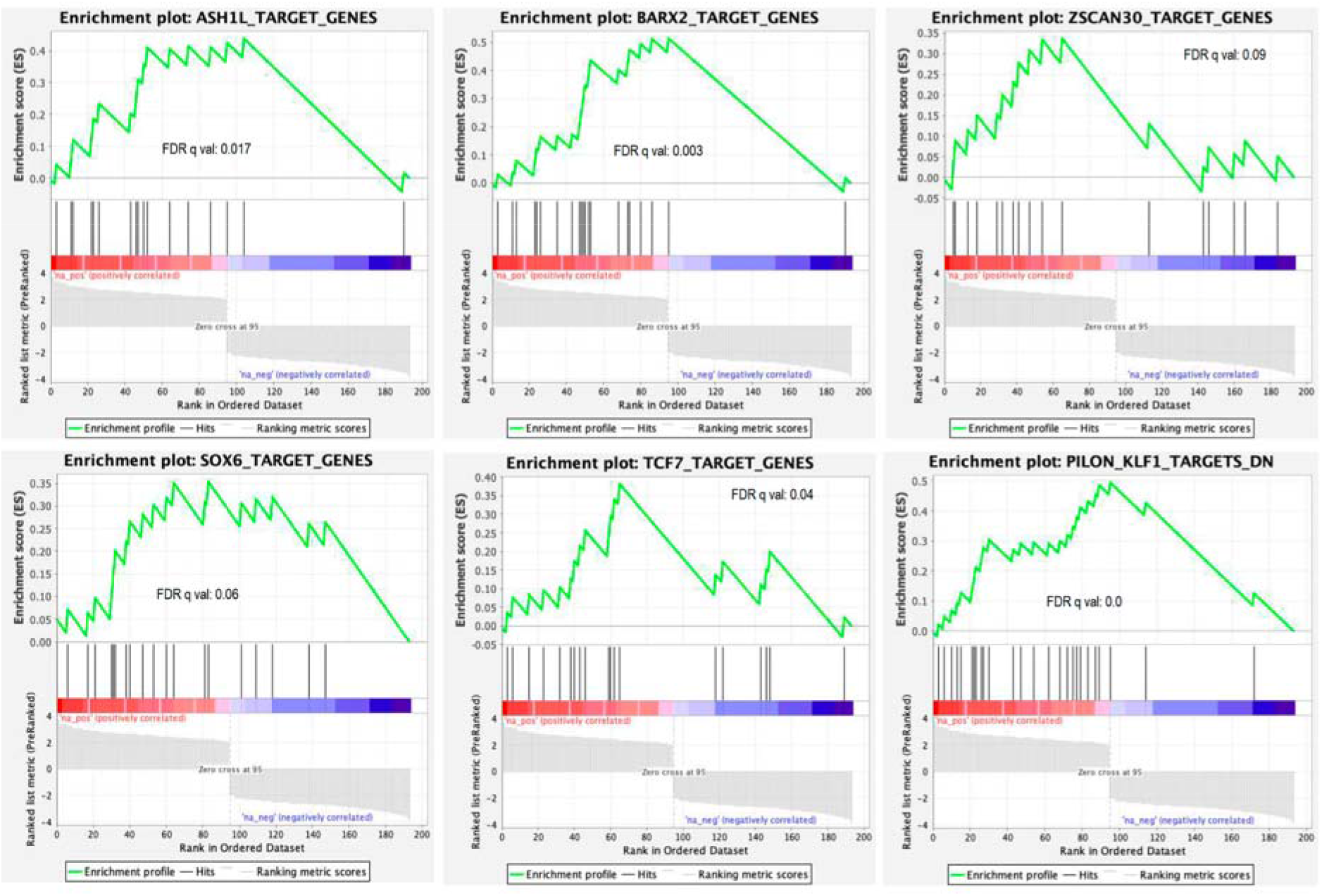
Gene set enrichment analysis (GSEA) of our gene set on the transcripts regulated by the factors ASH1 L, BARX2, ZSCAN30, SOX6, TCF7 and KLF.

Altogether, these data suggest a link between our primary gene set and the regulation of proliferation and pluripotency, suggesting that at least some of the genes in the list may be related to tumorigenesis and stemness in oral cavity tumors.

### 2.3 The identified gene sets stratify patients based on overall survival

Next, we analyzed the ability of these genes to be prognostic markers of OCSCC. We used the SurvExpress portal to investigate the potential prognostic value of the signature, including the genes of both the ‘long list’ and the ‘short list’ on different datasets as validation. We analyzed the prognostic value of the gene sets for OS (TCGA 2016 HNSCC dataset), cancer-free survival (oral cavity GSE26549 dataset), and metastasis-free survival (HNC tumors from the E-MTAB-1326 database) (Figure 5). We observed that the Cox regression curves showed statistically significant stratifications between high- and low-risk patients in the three databases. In HNC tumors, the risk group hazard ratio was 3.73 with a confidence interval from 2.48 to 5.63 (p=3.11e-10), providing a very good stratification of patients, and the Kaplan-Meier curve showed that the 50% OS of low-risk patients was approximately 100 months, while that of the high-risk group was below 25 months (Figure 5A). The signature of the ‘short list’ of genes still provides a very good stratification of patients, with a lower risk group hazard ratio (2.21) but still highly statistically significant (p=1.5e-07) (Figure 5B).

**Figure 5.**
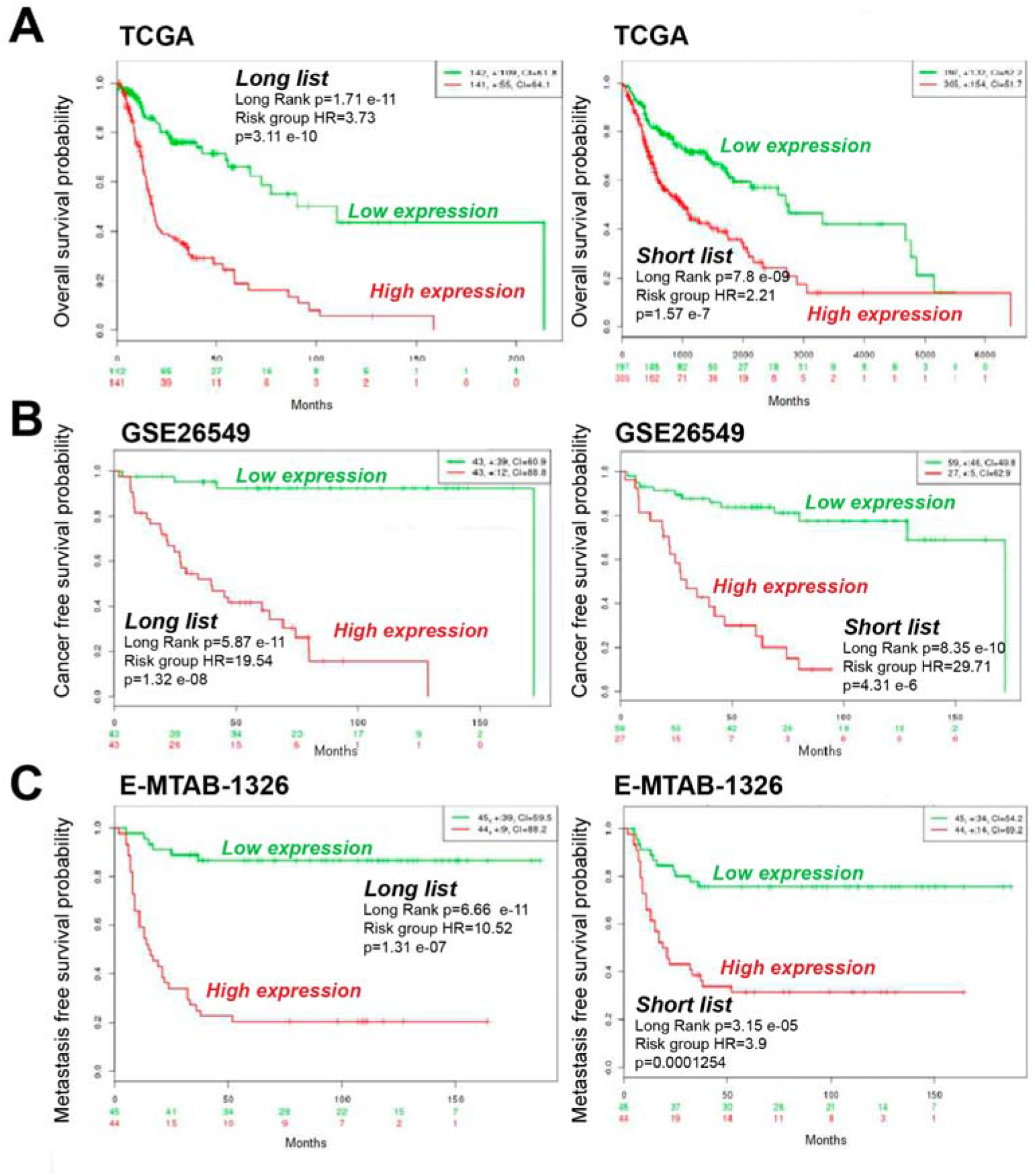
The identified gene sets stratify patients based on overall survival. (A) Kaplan-Meier plots showing the overall survival of patients with high (red) or low (green) expression levels of short- and long-list genes in the TCGA patient cohort databases. (B) Kaplan-Meier plots showing the cancer-free survival of patients with high (red) or low (green) expression levels of short- and long-list genes in the GSE26549 patient cohort database. (C) Kaplan-Meier plots showing metastasis-free survival of patients with high (red) or low (green) expression levels of short- and long-list genes in the E-MTAB-1328 patient cohort database. Data were analyzed with the log-rank test, and the associated P-values are shown in the graphs.

Analysis of the ‘long list’ signature in the oral cavity GSE26549 dataset for cancer-free survival showed a risk group hazard ratio of 19.54 with a confidence interval from 5.86 to approximately 65.17 (p=1.3e-6). Low-risk patients have a survival of approximately 90% beyond 150 months, while high-risk patients had decreased survival with a 50% survival below 4 months (Figure 5C). The ‘short list’ showed a risk group hazard ratio of 29.71 and was still statistically significant (p=4.31e-6) (Figure 5D). Similarly, analysis of the patient stratification on metastatic-free survival probability showed a risk group hazard ratio of 10.52 and statistical significance (p=1.31e-07) for the ‘long list’ (Figure 5E) and a risk group hazard ratio of 3.9 and statistical significance (p=0.00012) for the ‘short list’ signature (Figure 5F). These data demonstrate the ability to stratify patients according to the prognostic probability of the identified signatures.

### 2.4 Validation of the signature and selection of candidate genes

The analyzed gene sets can be used to generate a small profile of genes to be tested according to the values provided for low or high risk. However, to select a low number of biomarkers suitable for application to clinical settings, we worked only with the ‘short list’ of genes. These genes were tested against a new cohort of 89 tumor samples collected from the Central University Hospital of Asturias (HUCA) (Supplementary Table 2). Each single tumor sample was tested for mRNA expression levels of each gene, and the data were blindly analyzed by statistical Cox regression against clinical overall survival using the SPSS statistical program. We obtained 3 genes, WDR66, SERPINH1 and ZNF622, with statistical significance or a trend to statistical significance for prognosis among 15 candidates (Table 1). These 3 genes, especially WDR66,could stratify patients with higher or lower OS according to their mRNA levels (Figure 6). Analysis of the potential interacting proteins showed apparent independence among the different proteins (Supplementary Figure S4).

**Figure 6.**
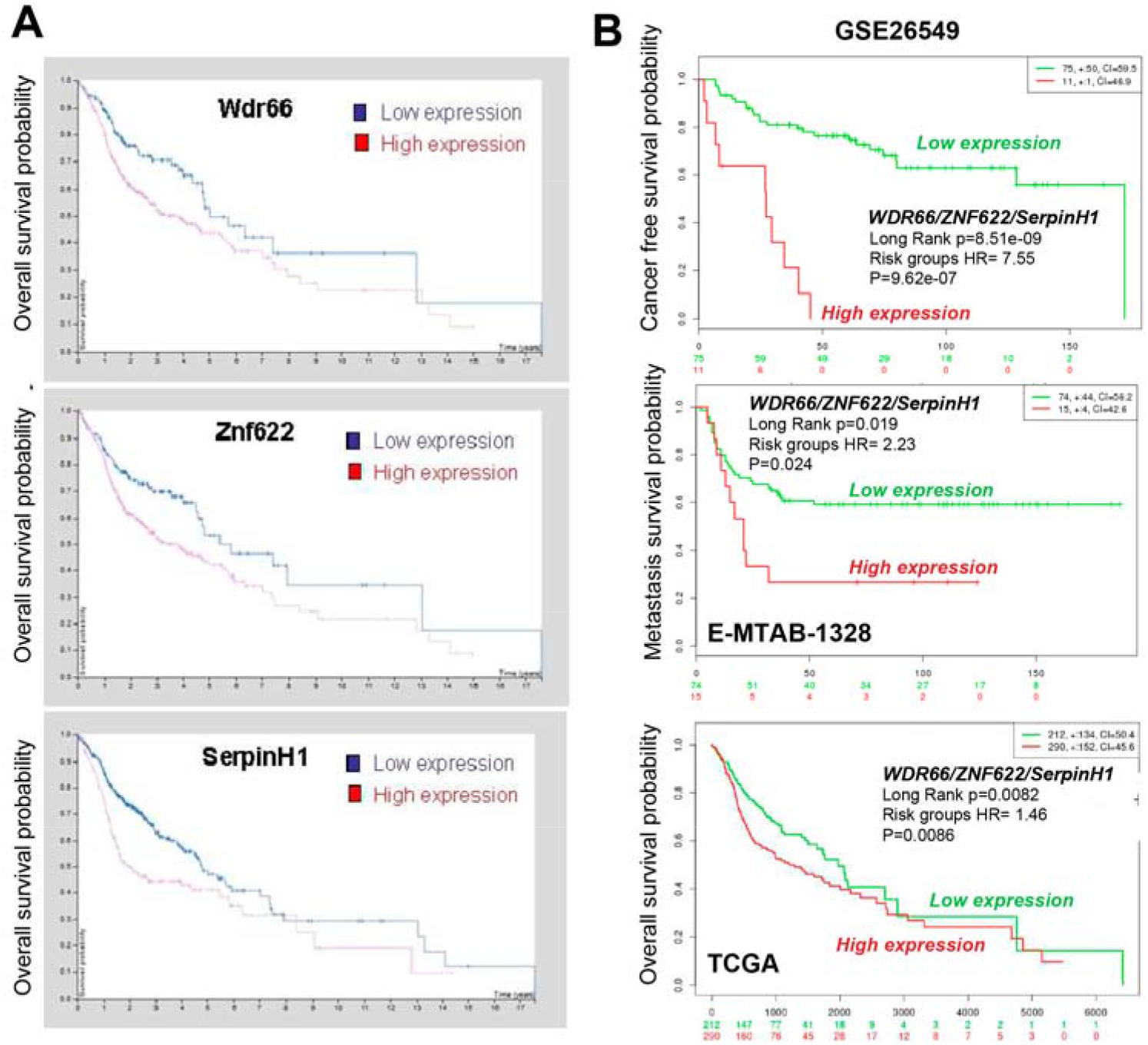
WRD66, SERPINH1 and ZNF622 genes stratify patients based on overall survival. (A) Kaplan-Meier plots showing the overall survival of patients with high (red) or low (blue) WRD66, SERPINH1 or ZNF622 expression levels in our Central University Hospital of Asturias (HUCA) patient cohort. Data were analyzed with the log-rank test, and the associated P-values are shown in the graphs. (B) Kaplan-Meier plots showing the overall survival of patients with high (red) or low (green) WRD66, SERPINH1 and ZNF622 expression levels in the GSE26549, E-MTAB-1328 and TCGA patient cohort databases. Data were analyzed with the log-rank test, and the associated P-values are shown in the graphs.

**Table 1.** List of 15 candidate genes obtained of cohort of 98 tumor samples collected from the Central University Hospital of Asturias (HUCA)

| Gene | Baseline | DFS<br>(Log Rank) | OS<br>(Log Rank) |
| --- | --- | --- | --- |
| *WDR66* | P75 | 0.033 | 0.003 |
| *ZNF622* | P25 | 0.123 | 0.086 |
| *SERPINH1* | P75 | 0.123 | 0.113 |
| *SYNGR1* | P25 | 0.186 | 0.167 |
| *ECT2* | Mean | 0.289 | 0.163 |
| *CPTML1* | P25 | 0.287 | 0.200 |
| *STAT1* | P75 | 0.265 | 0.224 |
| *BOC* | P75 | 0.719 | 0.232 |
| *FNCD3* | Mean | 0.248 | 0.262 |
| *ALSC2* | P25 | 0.246 | 0.330 |
| *PSMB2* | P25 | 0.486 | 0.337 |
| *LOC441178* | P75 | 0.419 | 0.358 |
| *CRNN* | Mean | 0.251 | 0.460 |
| *PPP13C* | P75 | 0.763 | 0.493 |
| *CGNL1* | P25 | 0.340 | 0.587 |

We have shown above that the expression levels of the genes WDR66, SERPINH1 and ZNF622 could individually stratify patients according to their OS at mRNA level. The cancer stem cell (CSC) phenotype of tumor cells is responsible for chemotherapy resistance and metastasis in HNSCC [26][27]; therefore, we tested the expression levels of these genes was cells enriched in CSCs

Based on the data obtained so far, we hypothesized that the SERPINH1, WDR66 and ZNF622 genes could be targets of pluripotency genes. To investigate this, we analyzed the presence of the DNA binding motifs of pluripotency transcription factors (TFs) near those genes. First, the genomic regions surrounding those genes were visualized in the WashU Epigenome Browser using high-resolution in situ Hi-C contact maps in hESCs [28]. Hi-C experiments use genome-wide chromosome conformation capture to analyze the 3D structure of chromatin, resulting in contact matrices represented as heatmaps whose intensity is indicative of the contact frequency of two different genomic regions, often located far away in the linear DNA sequence. Visual inspection of Hi-C data led to the identification of the topologically associated domain (TAD, identified by eye as red ‘triangles’) containing each gene, which constitutes its putative regulatory landscape (RL). Regulatory elements falling inside the RLs were determined using DNase-seq in hESCs (GSM736582), which indicates DNase hypersensitive sites (DHSs), i.e., open chromatin regions with bound transcription factors (TFs) that are more accessible for DNase I nuclease activity.

Motif enrichment analyses using Homer software did not show enrichment of pluripotency TF motifs inside the analyzed DHSs compared to genome-wide DHSs. However, several motifs associated with the pluripotency TFs Sox2, Oct4, Nanog, c-Myc and Klf4 were found. DHSs with more than 2 different motifs of pluripotency TFs were found for the three genes and are highlighted in Figure 7. Therefore, these results suggest that pluripotency TFs may contribute to SERPINH1, WDR66 and ZNF622 gene expression regulation. However, there is no evidence that these motifs are functional and bound by pluripotency factors in HNSCC. To test this point, we overexpressed these transcription SOX2 and OCT4 factors in HEK293 cells for 48 hours and measured the transcriptional variations of the SERPINH1, WDR66 and ZNF622 genes. We observed that the expression of ectopic SOX2 or CMYC in these primary cells triggers transcriptional activation of the 3 genes, SERPINH1, WDR66 and ZNF622, with an increase in their mRNA levels (Figure 8), confirming that these genes are targets of SOX2 or CMYC genes. Together, these data indicate that SERPINH1, WDR66 and ZNF622 are linked to the CSC phenotype in HNSCC and are likely direct targets of pluripotency TFs.

**Figure 7.**
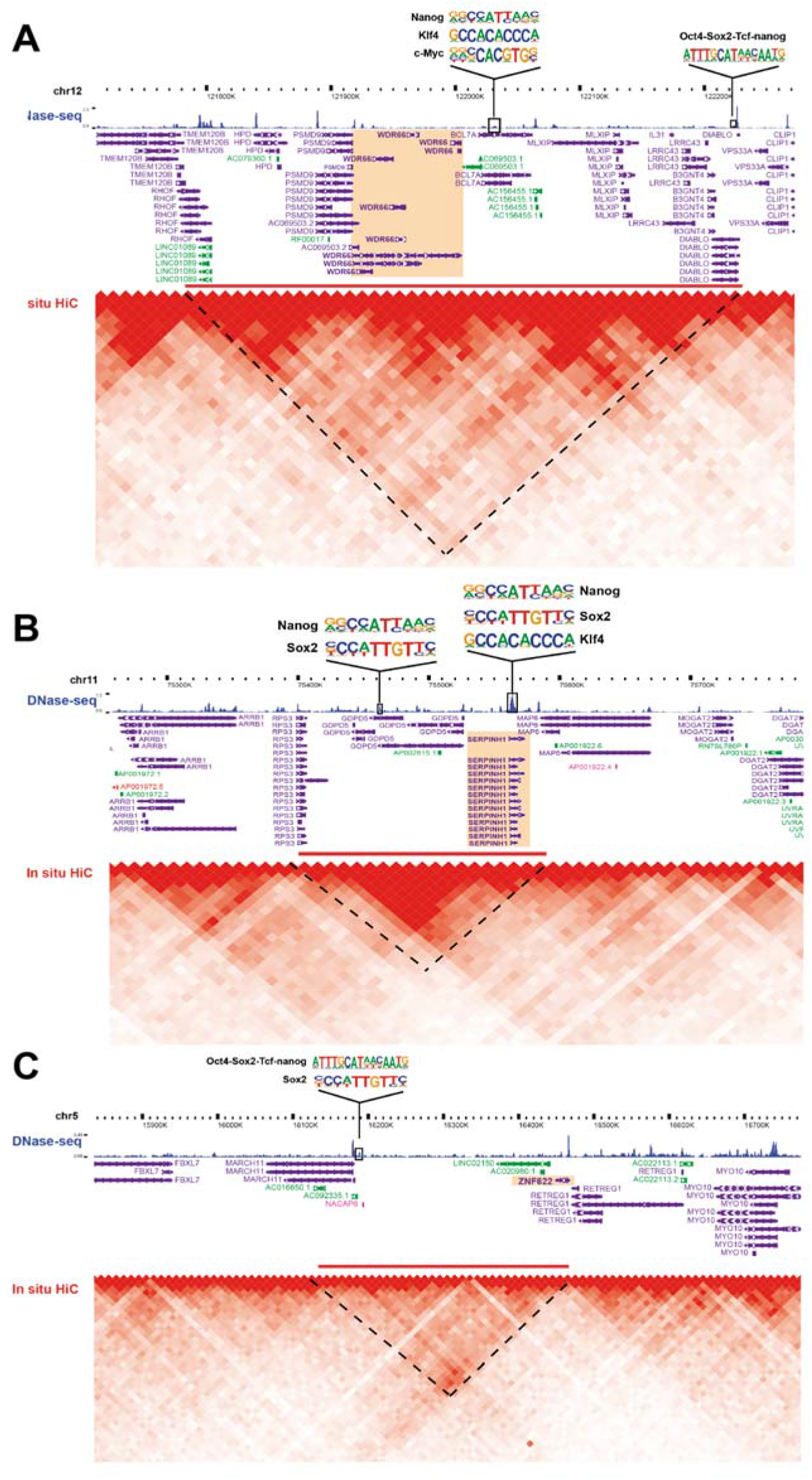
Pluripotency factor motifs in putative gene regulatory elements of WRD66, SERPINH1 and ZNF622 genes. Screenshots from the WashU Epigenome Browser showing the genomic region surrounding the A) WRD66, (B) SERPINH1 and (C) ZNF622 genes. Gene annotation (Genecode V29), in situ Hi-C contact maps (Krietenstein et al., bioRxiv 2019) and DNase-seq with DNase hypersensitive sites (GSM736582) in hESCs are shown. The WRD66, SERPINEH1 and ZNF622 genes are highlighted in orange, and their putative gene regulatory landscape based on Hi-C contacts is represented as a red bar. DNase hypersensitive sites inside this landscape containing predicted DNA binding motifs for several pluripotency factors are inside a black square, and the corresponding motif logos from the Homer motif database are shown on top.

**Figure 8.**
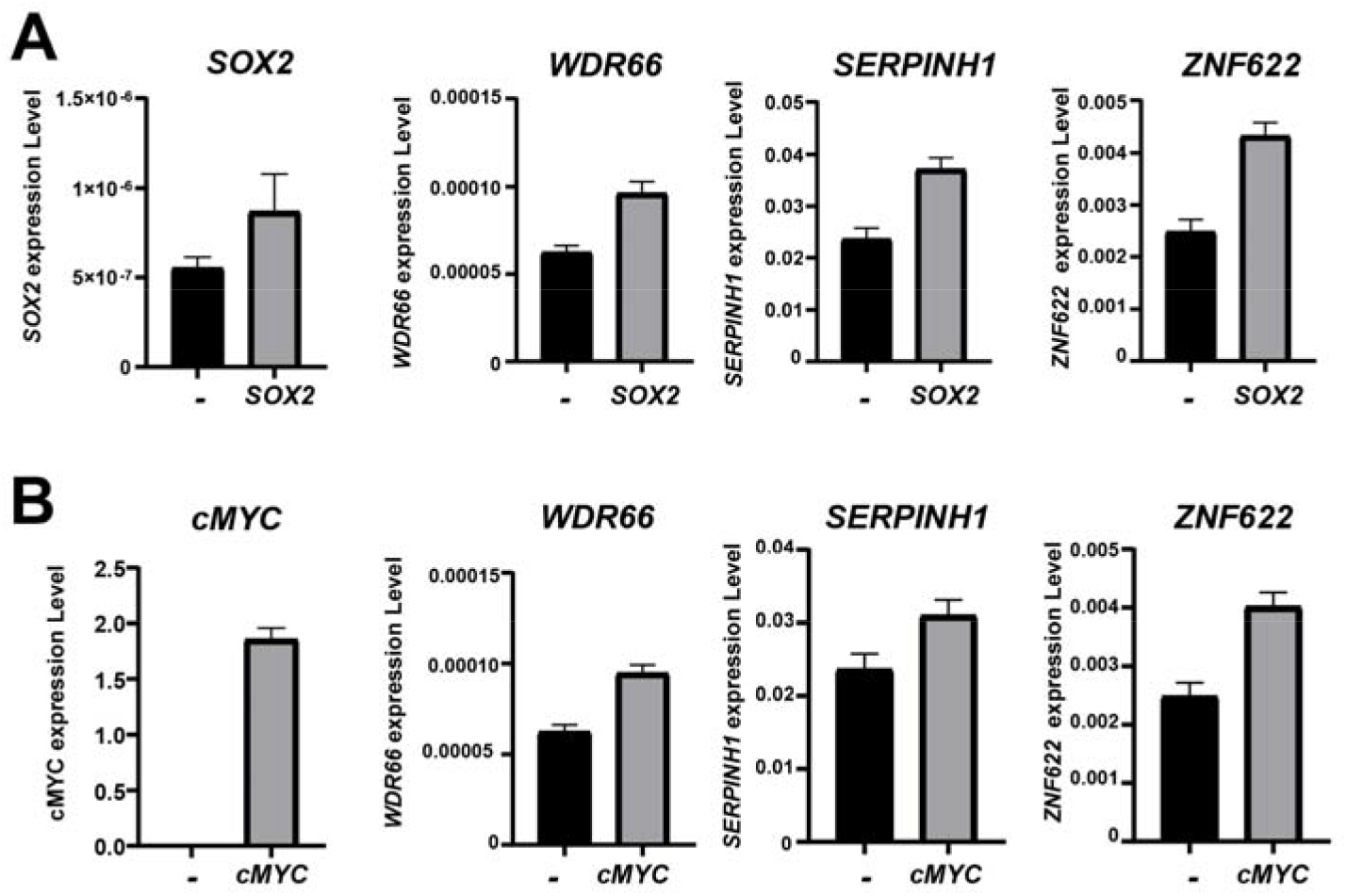
**(A) Analysis of the SOX2, WRD66, SERPINH1 and ZNF622 expression levels** by RT-qPCR in HEK293T cell lines expressing an EV (black) or overexpressing SOX2 (gray). (B) Analysis of the CMYC, WRD66, SERPINH1 and ZNF622 expression levels by RT-qPCR in HEK293T cell lines expressing an EV (black) or overexpressing CMYC (gray).

## 3. DISCUSION

In this study, we started with a bioinformatic analysis of oral cancer databases to identify a list of genes with clear diagnostic capability to separate tumor and nontumor samples according to their expression profiles. These genes belong to different functional pathways but seem to be more related to the cell cycle, inflammation and stemness. According to the Cox analysis, this profile clearly stratified: i) patients with oral cancers, and ii) those with better and worse prognosis in total HNSCC tumors. Validation of this list of genes in an independent cohort reduced it to 3 representative genes: WDR66, ZNF622 and SERPINH1, which together can establish the diagnostic and prognostic value of the genetic signature. Interestingly, these genes correlate with the EMT and stemness capabilities of cells and are transcriptional targets of the pluripotency TFs Sox2 and c-Myc.

In a TCGA studies [22, 24], HNSCCs analyzed, in HPV-positive cases showed frequent loss of TRAF3, PIK3CA activating mutations and E2F1 amplifications, whereas the most common genetic events for HPV negative cases were co-amplifications of 11q13 and 11q22, focal deletions in tumor suppressor genes and focal amplifications in tyrosine kinase receptor genes.

Many mutations are responsible for resistance to anticancer drugs, especially platinum-based chemotherapy and radiotherapy, as used for many HNSCC tumors [16]. This genetic variability and complexity make the analysis of HNSCC cancer resistance considerably complex. We believe that the known mutations that confer therapy resistance are distributed among different pathways, which may activate a few common essential effector genes. Ultimately, these effector genes may be responsible for the HNSCC cancer resistance physiology, and this physiology may be reflected in a few biomarkers, which may be measurable, predictive and targetable. Therefore, genetic events that alter cell physiology may limit the efficacy of therapies with anticancer compounds. Many mutations or alterations of epigenetic signals and methylation profiles are involved in therapy resistance (TCGA database, cBioPortal). In addition, a recent analysis of patients with HNSCC tumors showed a high complexity, a high number of genomic and genetic alterations, and high levels of intra- and intertumoral heterogeneity [16, 22, 24]. Hence, the use of these alterations, most of them occurring at low frequencies, as biomarkers to predict sensitivity or resistance is not currently useful, in HNSCC, especially considering the large number of HNSCC tumor types. Because resistance to treatments is one of the main causes of poor survival among HNSCC cancer patients, the identification of prognostic and predictive biomarkers, as well as understanding the mechanisms driving resistance, is urgently needed.

Using 11 public databases with HNSCC patient data, we identified the most common and strong DEGs, obtaining two lists of genes covering the top 1% and top 5% of those genes. By analyzing the expression profiles of those genes in the selected databases, we found a clear separation of tumoral and nontumoral samples with distinct expression profiles of genes for both the long and the short lists. GSEA indicated that cellular functions related to the cell cycle, the DNA damage response, cytokine signaling, and the immune system were enriched among the identified lists, they were also enriched in the embryonic stem cell core, as well as in target genes of several pluripotency regulators, providing a first link between these genes and tumor stemness. Interestingly, both the long and short lists could separate low- and high-risk groups of patients according to their OS with statistical significance, indicating that they were indeed associated with the survival of patients. These lists of genes constitute good genetic signatures that predict patient outcome. However, they are composed of many genes, and we aimed to reduce the list to a few representative genes using a validation cohort. This analysis reduced the signature to three genes that were not functionally related but could separate patients by their differential OS rate: WDR66, SERPINH1 and ZNF622.

WD repeat-containing 66 gene (WDR66 located on chromosome 12 (12q24.31), belongs to the WD-repeat protein family, a large family found in all eukaryotes that is involved in various functions ranging from signal transduction and transcriptional regulation to cell cycle control, autophagy and apoptosis [29]. These repeating units are believed to serve as scaffolds for multiple protein interactions with various proteins. Mutations in WDR66 have been linked to male infertility [30][31], and in cancer, high WDR66 expression has been associated with poor OS of patients suffering from esophageal squamous carcinomas [32]. SERPINH1 codes a serine-type endopeptidase inhibitor, also known as collagen binding protein 1 or heat shock protein 47. The SERPINH1 protein is involved in tumor angiogenesis, growth, migration and metastasis in several types of cancer [33], particularly HNSCC [34]. Finally, ZNF622 is a zinc finger protein overexpressed in colorectal tumors [35, 36] that is associated with the assembly of the ribosomal 60S subunit. In this regard, ZNF622 has been recently linked to hematopoietic stem cell regeneration and is a target of the E3 ubiquitin ligase HectD1, which controls this process by controlling ribosomal assembly [37].

We demonstrated that the three genes composing the final signature are differentially expressed in HNSCC tumors and in HNSCC cell lines, indicating that they are indeed deregulated in CSCs. Interestingly, using public Hi-C and DNase-seq data, we found several instances of DNA binding motifs for pluripotency TFs within open chromatin regions that could potentially control the expression of these genes. Confirmation of this regulation was demonstrated in overexpression experiments of SOX2 and CMYC pluripotency genes, in which we detected significant increases in the expression of the three candidate genes (Figure 8). Together, these data link WDR66, SERPINH1 and ZNF622 to the CSC phenotype and stemness in HNSCC tumors, suggesting that they might be effectors of stemness in tumor cells and common markers of worse prognosis and therapy resistance

In conclusion, the genetic profile identified in this work could be used to stratify patients due to its predictive value for therapy resistance, but it would also be interesting to combine our profile with other clinical predictors, such as tumor stage, histological type, or the degree of differentiation, to provide an accurate clinical assessment with increased prognostic and predictive value that could help stratify patients in clinical practice.

## 4. MATERIALS AND METHODS

### Cell Culture

The established cell lines RPMI-2650 and Detroit-562 cells were obtained from the ATCC commercial repository. Cells were maintained in DMEM (Sigma) supplemented with 10% fetal bovine serum (FBS) (Gibco), penicillin, streptomycin and fungizone (Sigma). For the biopsy-derived cell lines, larynx primary cell lines were obtained from the Biomedical Research in Cancer Stem Cells Group of Barcelona. Cells were maintained in RPMI 1640 (Sigma) supplemented with 20% FBS, 1% glutamine, penicillin, streptomycin and fungizone as described [38]. All cell lines were negative for mycoplasma.

### Tumorsphere assay

Ten thousand cells of RPMI-2650 and Detroit celllines were seeded in 24-well Ul-tra-Low Attachment Plates (Costar) containing 1 mL of MammoCult basal medium (Stem Cell Technologies) supplied with 10% MammoCult proliferation supplement, 6 μg/mL heparin, 0.48 μg/mL hydrocortisone, penicillin and streptomycin. After 1 week, the number and size of tumorspheres formed in three wells were measured using an inverted microscope (Olympus IX-71) and cellSens Dimension software. All of the tumorspheres from the plate were harvested for subsequent RNA extraction.

### RT-qPCR

Total RNA from cell lines and tumorspheres was extracted and purified using the RNeasy Mini Kit (QIAGEN). Reverse transcription was performed with 1 μg of RPMI-2650 and Detroit-562 mRNA or 0.221μg of primary cell line mRNA using the High Capacity cDNA Reverse Transcription kit (Life Technologies). The qPCR mixture (10 μL) contained 2 μL of cDNA (diluted 1:3 in the case of RPMI-2650 and Detroit-5622.5 μL of water, 5 μL of GoTaq^®^ Probe qPCR Master Mix (Promega) and 0.5 μL of the appropriate TaqMan Assay (20X) (Applied Biosystems). We used the following probes: GAPDH (Hs03929097_g1) as an endogenous control, WRD66 (Hs00376225_m1), SERPINH1 (Hs01060397_g1), ZNF622 (Hs003691082_m1), SOX2 (Hs01053049_s1), CMYC (Hs00153408_m1).

### Study approval

Sample use and experimental procedures were performed in accordance with the Declaration of Helsinki. Written informed consent was obtained from all patients. Formalin-fixed paraffin-embedded (FFPE) tumor biopsies and data from donors were provided by the Principado de Asturias BioBank (PT17/0015/0023), integrated in the Spanish National Biobanks Network, and histological diagnosis was confirmed by an experienced pathologist. Samples were processed following standard operating procedures with the appropriate approval of the Ethical and Scientific Committees of the Hospital Universitario Central de Asturias and the Regional CEIm from Principado de Asturias (date of approval May 14th, 2019; approval number: 141/19, for the project PI19/00560) Also, Written informed consent was provided by all patients. This project was approved by the Research Ethics Committee of the Hospital Universitario Virgen del Rocio (CEI 0309-N-15).

### Patient cohort

Surgical tissue specimens from 89 patients with HPV-negative HNSCC who underwent resection of their tumors at the Hospital Universitario Central de Asturias between 2000 and 2009 were retrospectively collected. The characteristics of the studied cases are shown in Supplementary Table 2. All patients had a single primary tumor, microscopically clear surgical margins and received no treatment prior to surgery. The stage of the tumors was determined according to the TNM system of the International Union Against Cancer (7th Edition). 68 (76%) of 89 patients received postoperative radiotherapy.

Analyses of cancer patient databases: To validate our results, we obtained data from publicly available clinical and genomic databases, including Oncomine (https://powertools.oncomine.com/) and the TCGA Research Network (http://cancergenome.nih.gov/). We performed meta-analyses of the PrognoScan public patient datasets (http://dna00.bio.kyutech.ac.jp/PrognoScan/) to analyze the expression levels in tumor and nontumor databases for HNSCC tissue samples. Statistical significance versus normal samples was considered to be P < 0.05. Patient survival was analyzed using the R2 Genomics analysis and visualization platform (http://hgserver1.amc.nl), developed by the Department of Oncogenomics of the Academic Medical Center (AMC) (Amsterdam, Netherlands). Kaplan-Meier plots showing patient survival were generated for databases with available survival data using the scan method, which searched for the optimum survival cutoff based on statistical analyses (log-rank test), thus finding the most significant expression cutoff. To analyze the protein network, we used the web portal https://string-db.org.

### Quantification and statistical analysis

All statistical analyses were performed using GraphPad Prism 4. The distribution of quantitative variables among different study groups was assessed using parametric (Student’s t-test) or nonparametric (Kruskal–Wallis or Mann–Whitney) tests, as appropriate. Experiments were performed a minimum of three times and were always performed as independent triplicates. Survival data from the patient databases were analyzed with the log-rank Mantel-Cox statistical test.

## DECLARATIONS

### Ethics approval and consent to participate

Sample use and experimental procedures were performed in accordance with the Declaration of Helsinki. Written informed consent was obtained from all patients. Samples were processed following standard operating procedures with the appropriate approval of the Ethical and Scientific Committees of the Hospital Universitario Central de Asturias and the Regional CEIm from Principado de Asturias (date of approval May 14th, 2019; approval number: 141/19, for the project PI19/00560)

## Acknowledgments

The authors thank the donors and the biobank of the Hospital Universitario Principal de Asturias y Virgen del Rocío-Instituto de Biomedicina de Sevilla (Andalusian Public Health System Biobank and ISCIII-Red de Biobancos PT17/0015/0041 and PT17/0015/0023) for the human specimens used in this study.

## Authors’contributions

LS-D, SM-G and AC designed the experiments; LS-D, SM-G, BF, MG-C, JD-P, ES_M,EMV-S, LEN, DO-A, JJM-L, MAC, JP-S, MPJ-G, JMG-H, MP, AG-Q, and AC, performed the experiments; MDC, JPR and PE-G collected patients, tissues and clinical data; YG-M and MLL generated primary cell lines from patients; SM-G, AC, JPR-T and AC analyzed and interpreted the data; SMG, JMG-H and AC wrote the manuscript. All authors read, edited and approved the final manuscript.

## Consent for publication

All authors read and approve the manuscript and give consent for publication.

---

**Supplementary table 1.**
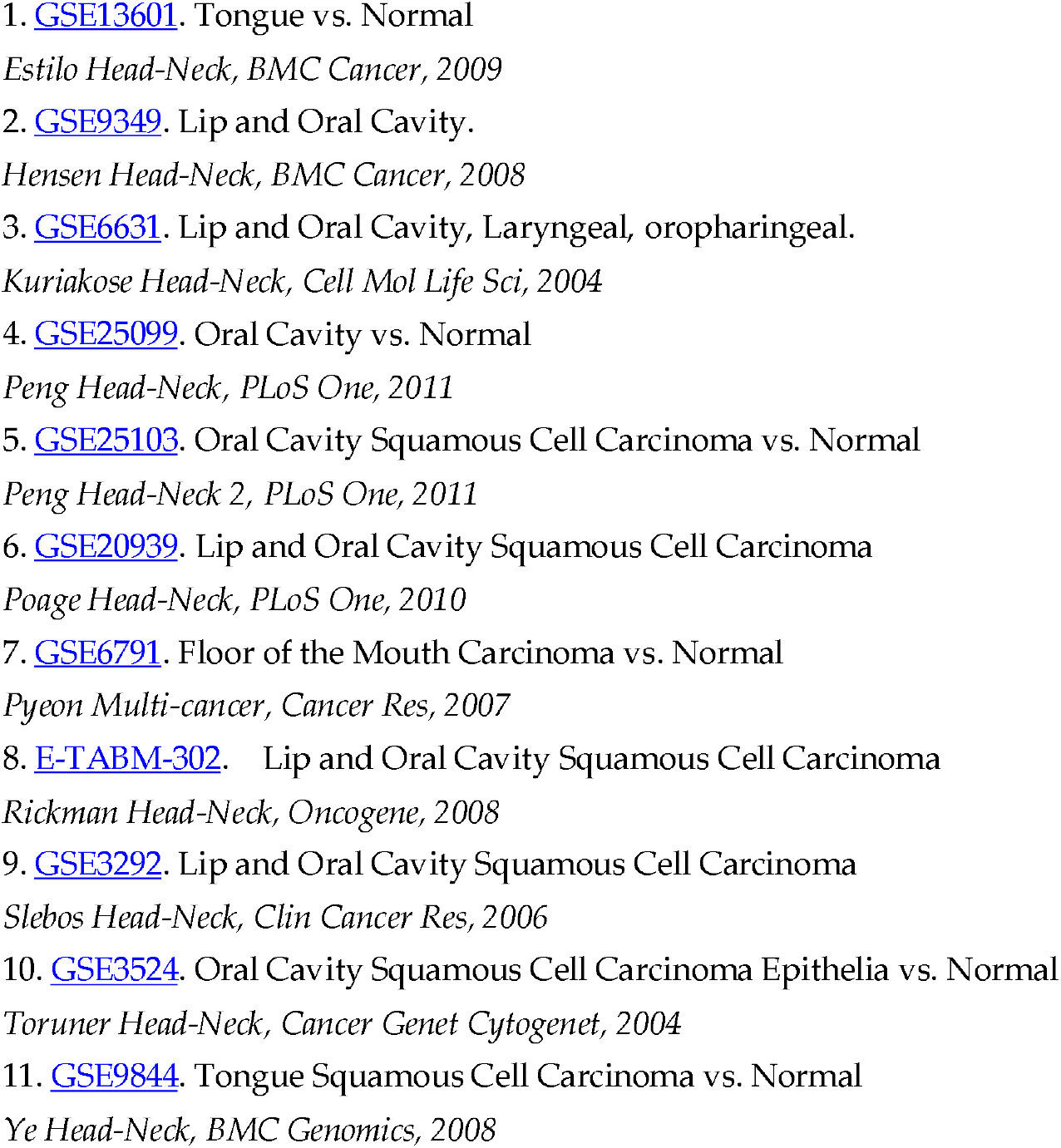
11 Public databases used in the screening for the figure 1 and table 1.

1. Laryngeal, Oral Cavity, Oropharyngeal, Tongue, pharyngeal, lips, Tonsillar and other tumors of epithelial origing. All in the oral cavity.

**Supplementary table 2.** Clinical data of the Hospital Universitario Central de Asturias.

Clinical data of the Hospital Universitario Central de Asturias
|  | Total number | % |
| --- | --- | --- |
| Nº patients | 89 |  |
| Radiotherapy | 68 | 76,40 |
| Location: tongue base | 38 | 42,70 |
| Location: tonsil | 51 | 57,30 |
| T1 | 8 | 8,99 |
| T2 | 13 | 14,61 |
| T3 | 24 | 26,97 |
| T4 | 44 | 49,44 |
| T1+T2 | 67 | 75,28 |
| T3+T4 | 22 | 24,72 |
| N0 | 8 | 8,99 |
| N+ | 81 | 91,01 |
| M1 | 0 | 0,00 |
| Grade 1 | 37 | 41,57 |
| Grade 2 | 30 | 33,71 |
| Grade 3 | 22 | 24,72 |
| Stage I | 0 | 0 |
| Stage II | 0 | 0 |
| Stage III | 7 | 7,87 |
| Stage IVA | 79 | 88,76 |
| Stage IVB | 3 | 3,37 |
| Stage IVC | 0 | 0 |
| No recurrence | 30 | 33,71 |
| Recurrence | 59 | 66,29 |
| Average free survival (months) | 6,53 | 7,34 |
| Average monitoring time (months) | 28,30 | 31,80 |
| Exitus | 69 | 77,53 |
| No exitus | 20 | 22,47 |
| Exitus tumor | 55 | 61,80 |
| No exitus tumor | 34 | 38,20 |

**Supplementary Figure 1.**
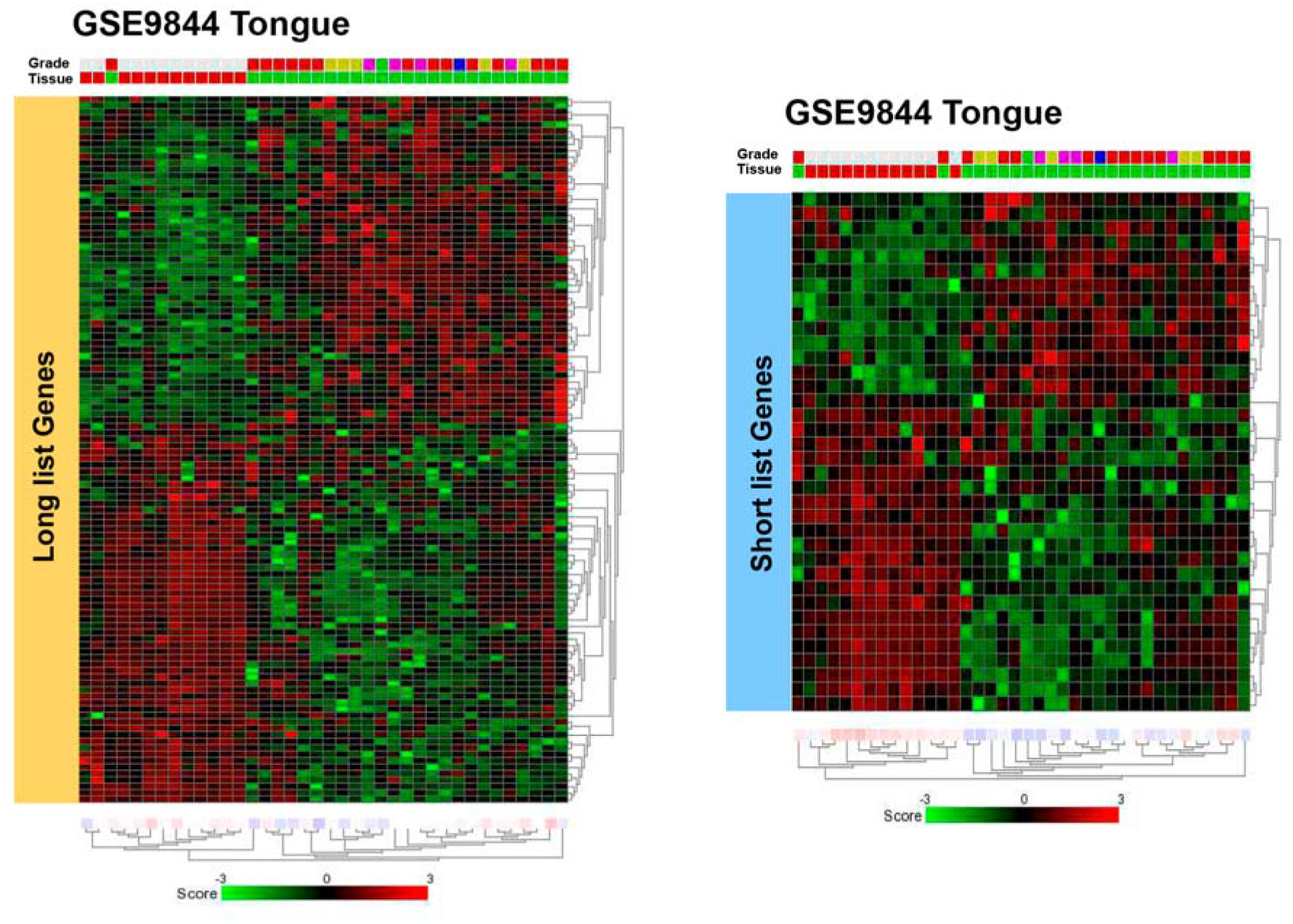
Both long and short gene sets transcriptional heat map of patients and control individuals in the tongue tumors GSE9844 database.

**Supplementary Figure 2.**
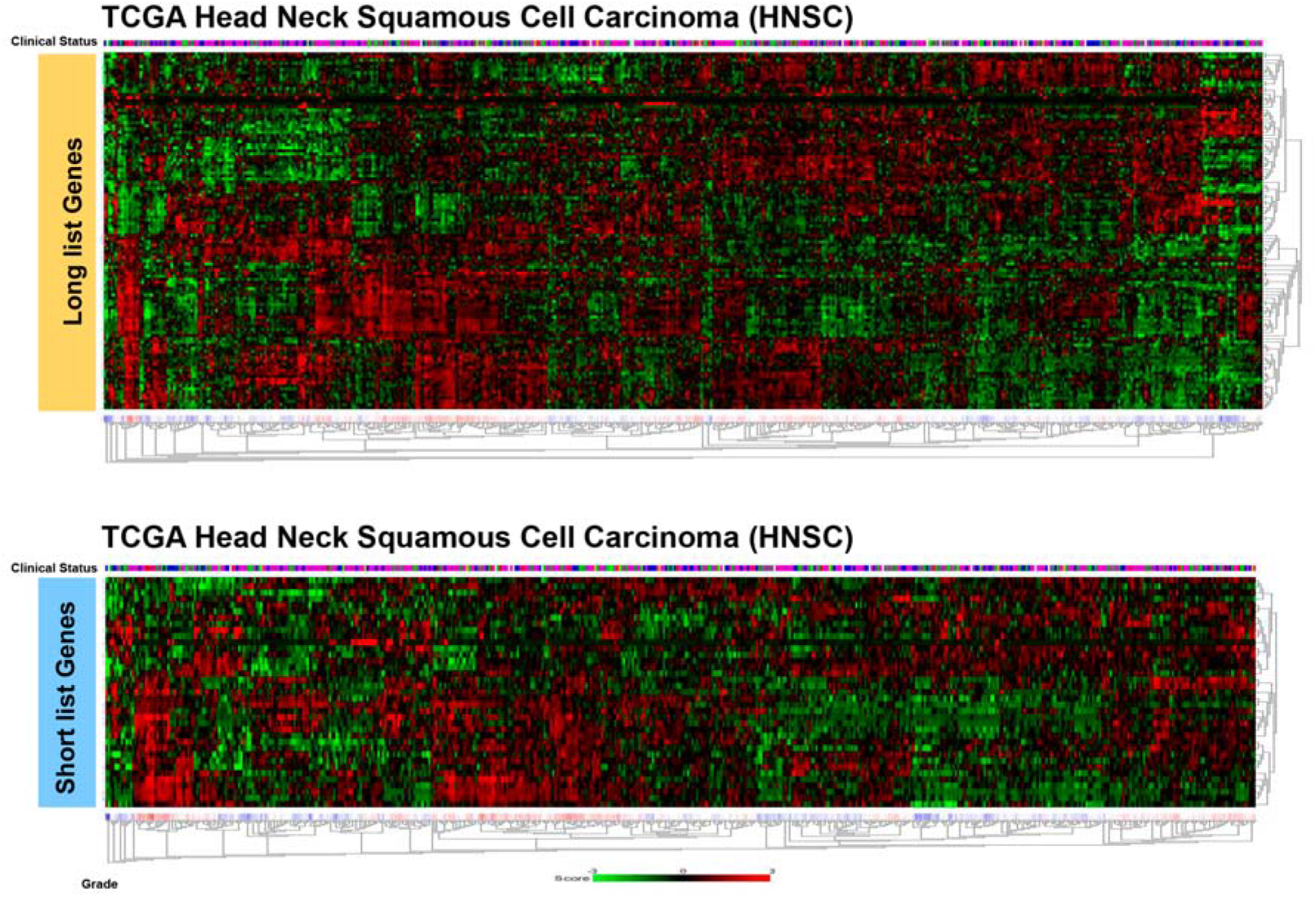
Both long and short gene sets transcriptional heat map of patients and control individuals in the HNSCC tumors of the TCGA database.

**Supplementary Figure 3.**
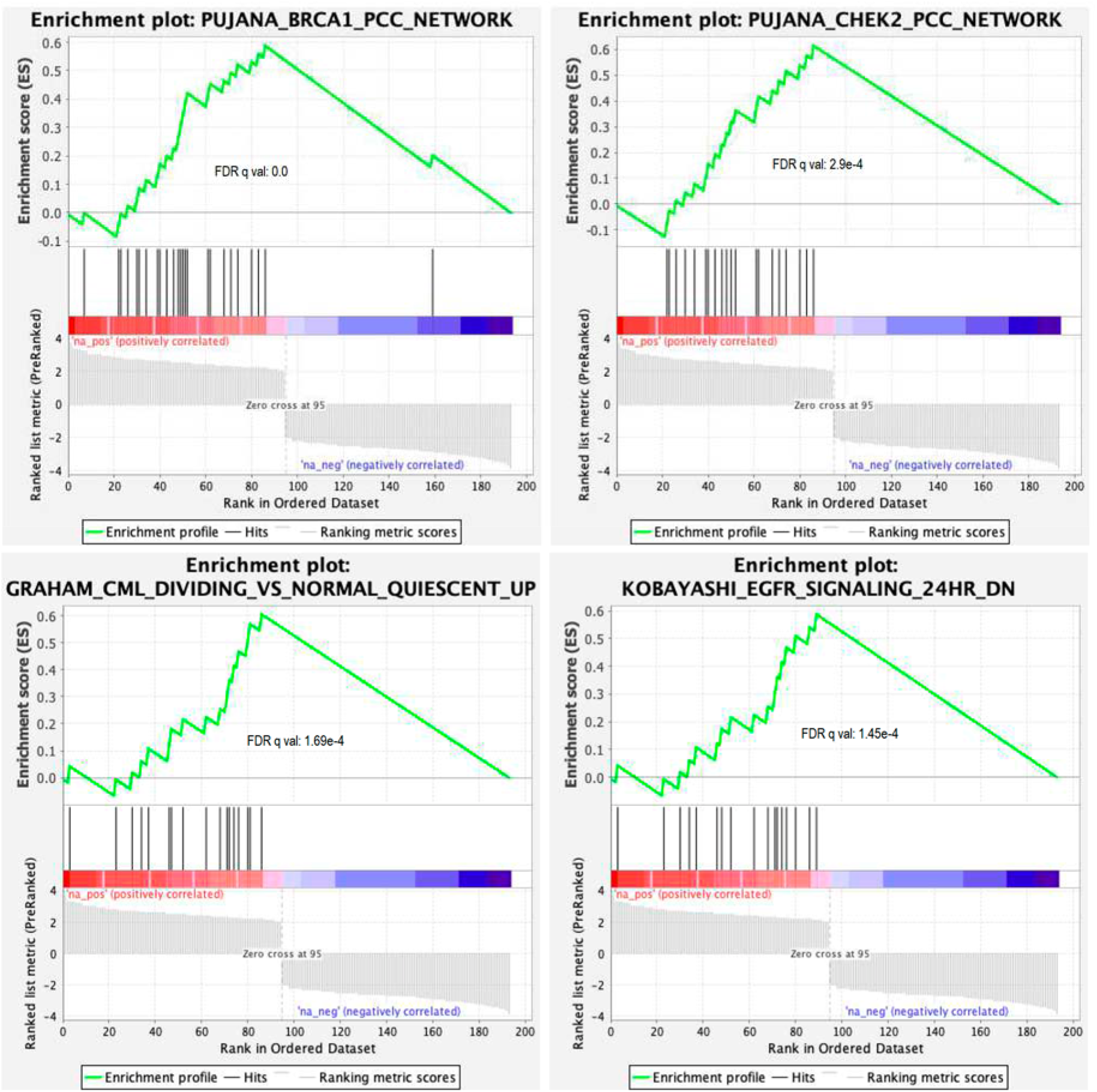
Network analysis of the gene sets. Gene set enrichment analysis (GSEA) correlations of the gene set with Gene Ontology (GO) terms in different specific datasets, with a bias towards the upregulated genes.

**Supplementary Figure 4.**
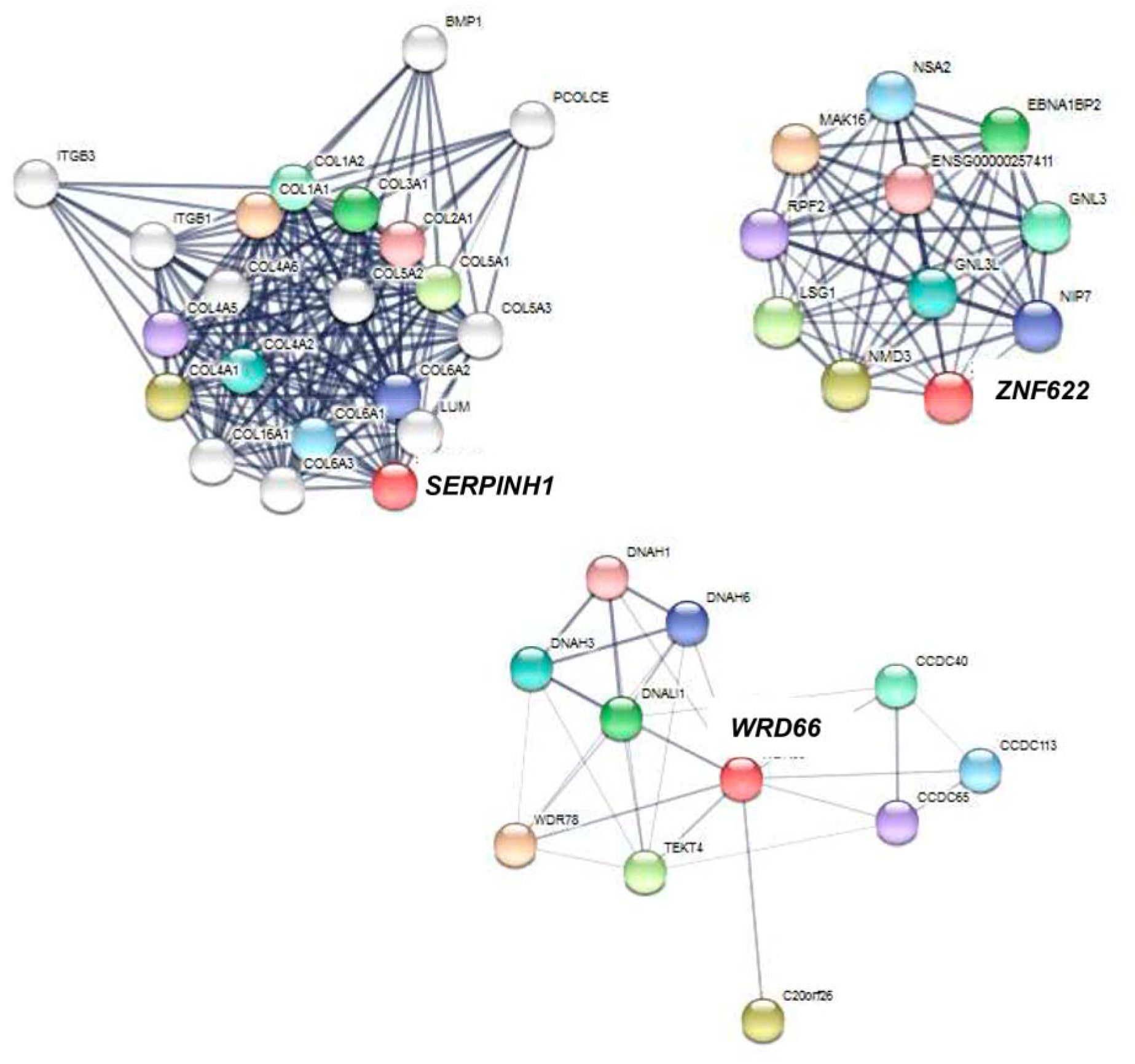
Protein interaction analysis of the SERPINH1, ZNF622 and WDR66 genes. Analysis was performed in STRING portal up to the 3rd interaction step.

## REFERENCES

1. Ferlay, J., et al., Estimates of worldwide burden of cancer in 2008: GLOBOCAN 2008. Int J Cancer, 2010. 127(12): p. 2893–917.

2. Jemal, A., et al., Global cancer statistics. CA Cancer J Clin, 2011. 61(2): p. 69–90.

3. Fearon, E.R. and B. Vogelstein, A genetic model for colorectal tumorigenesis. Cell, 1990. 61(5): p. 759–67.

4. Bosatra, A., R. Bussani, and F. Silvestri, From epithelial dysplasia to squamous carcinoma in the head and neck region: an epidemiological assessment. Acta Otolaryngol Suppl, 1997. 527: p. 47–8.

5. Sinha, S. and A. Gajra, Nasopharyngeal Cancer, in StatPearls. 2021: Treasure Island (FL).

6. Goodson, W.H., 3rd, et al., Assessing the carcinogenic potential of low-dose exposures to chemical mixtures in the environment: the challenge ahead. Carcinogenesis, 2015. 36 Suppl 1: p. S254–96.

7. Venugopal, R., et al., Familial Cancers of Head and Neck Region. J Clin Diagn Res, 2017. 11(6): p. ZE01–ZE06.

8. Ryan, W.R., et al., Oncologic outcomes of human papillomavirus-associated oropharynx carcinoma treated with surgery alone: A 12-institution study of 344 patients. Cancer, 2021.

9. Lorenzatti Hiles, G., et al., Understanding the impact of high-risk human papillomavirus on oropharyngeal squamous cell carcinomas in Taiwan: A retrospective cohort study. PLoS One, 2021. 16(4): p. e0250530.

10. Partridge, M., et al., Field cancerisation of the oral cavity: comparison of the spectrum of molecular alterations in cases presenting with both dysplastic and malignant lesions. Oral Oncol, 1997. 33(5): p. 332–7.

11. de Miguel-Luken, M.J., M. Chaves-Conde, and A. Carnero, A genetic view of laryngeal cancer heterogeneity. Cell Cycle, 2016. 15(9): p. 1202–12.

12. Hussein, A. A., et al., A review of the most promising biomarkers for early diagnosis and prognosis prediction of tongue squamous cell carcinoma. Br J Cancer, 2018. 119(6): p. 724–736.

13. Fung, N., et al., The role of human papillomavirus on the prognosis and treatment of oropharyngeal carcinoma. Cancer Metastasis Rev, 2017. 36(3): p. 449–461.

14. Weiss, J. and D.N. Hayes, Classifying squamous cell carcinoma of the head and neck: prognosis, prediction and implications for therapy. Expert Rev Anticancer Ther, 2014. 14(2): p. 229–36.

15. Chow, L.Q.M., Head and Neck Cancer. N Engl J Med, 2020. 382(1): p. 60–72.

16. Loyo, M., et al., Lessons learned from next-generation sequencing in head and neck cancer. Head Neck, 2013. 35(3): p. 454–63.

17. Ferris, R.L. and L. Licitra, PD-1 immunotherapy for recurrent or metastatic HNSCC. Lancet, 2019. 394(10212): p. 1882–1884.

18. Hanahan, D. and R.A. Weinberg, The hallmarks of cancer. Cell, 2000. 100(1): p. 57–70.

19. Califano, J., et al., Genetic progression model for head and neck cancer: implications for field cancerization. Cancer Res, 1996. 56(11): p. 2488–92.

20. Grimminger, C.M. and P.V. Danenberg, Update of prognostic and predictive biomarkers in oropharyngeal squamous cell carcinoma: a review. Eur Arch Otorhinolaryngol, 2011. 268(1): p. 5–16.

21. Bockmuhl, U., et al., Genomic alterations associated with malignancy in head and neck cancer. Head Neck, 1998. 20(2): p. 145–51.

22. Ran, Q.C., et al., Mining TCGA database for prognostic genes in head and neck squamous cell carcinoma microenvironment. J Dent Sci, 2021. 16(2): p. 661–667.

23. Stransky, N., et al., The mutational landscape of head and neck squamous cell carcinoma. Science, 2011. 333(6046): p. 1157–60.

24. Jin, Y. and Y. Yang, Identification and analysis of genes associated with head and neck squamous cell carcinoma by integrated bioinformatics methods. Mol Genet Genomic Med, 2019. 7(8): p. e857.

25. Forastiere, A., et al., Head and neck cancer. N Engl J Med, 2001. 345(26): p. 1890–900.

26. Tahmasebi, E., et al., The current markers of cancer stem cell in oral cancers. Life Sci, 2020. 249: p. 117483.

27. Garcia-Mayea, Y., et al., Insights into new mechanisms and models of cancer stem cell multidrug resistance. Semin Cancer Biol, 2020. 60: p. 166–180.

28. Krietenstein, N., et al., Ultrastructural Details of Mammalian Chromosome Architecture. Mol Cell, 2020. 78(3): p. 554–565 e7.

29. Li, D. and R. Roberts, WD-repeat proteins: structure characteristics, biological function, and their involvement in human diseases. Cell Mol Life Sci, 2001. 58(14): p. 2085–97.

30. Kherraf, Z.E., et al., A Homozygous Ancestral SVA-Insertion-Mediated Deletion in WDR66 Induces Multiple Morphological Abnormalities of the Sperm Flagellum and Male Infertility. Am J Hum Genet, 2018. 103(3): p. 400–412.

31. Kherraf, Z.E., et al., Creation of knock out and knock in mice by CRISPR/Cas9 to validate candidate genes for human male infertility, interest, difficulties and feasibility. Mol Cell Endocrinol, 2018. 468: p. 70–80.

32. Wang, Q., C. Ma, and W. Kemmner, Wdr66 is a novel marker for risk stratification and involved in epithelial-mesenchymal transition of esophageal squamous cell carcinoma. BMC Cancer, 2013. 13: p. 137.

33. Duarte, B.D.P. and D. Bonatto, The heat shock protein 47 as a potential biomarker and a therapeutic agent in cancer research. J Cancer Res Clin Oncol, 2018. 144(12): p. 2319–2328.

34. Fang, X., et al., Long non-coding RNA SNHG22 facilitates the malignant phenotypes in triple-negative breast cancer via sponging miR-324-3p and upregulating SUDS3. Cancer Cell Int, 2020. 20: p. 252.

35. Mun, K. and T. Punga, Cellular Zinc Finger Protein 622 Hinders Human Adenovirus Lytic Growth and Limits Binding of the Viral pVII Protein to Virus DNA. J Virol, 2019. 93(3).

36. Kleivi, K., et al., Gene expression profiles of primary colorectal carcinomas, liver metastases, and carcinomatoses. Mol Cancer, 2007. 6: p. 2.

37. Lv, K., et al., HectD1 controls hematopoietic stem cell regeneration by coordinating ribosome assembly and protein synthesis. Cell Stem Cell, 2021.

38. Garcia-Mayea, Y., et al., TSPAN1: A Novel Protein Involved in Head and Neck Squamous Cell Carcinoma Chemoresistance. Cancers (Basel), 2020. 12(11).

